# Identification of *Acinetobacter baumannii* and detection of ß - lactam antibiotic resistance genes in clinical samples by multiplex PCR

**DOI:** 10.1101/2020.10.25.353896

**Authors:** Trinh Phan-Canh, Thao Le-Thi-Thanh, Thuy Ngo-Thi-Bich, Thanh Nguyen-Thi-Thanh, Linh Ho-Le-Truc, Tu-Anh Nguyen

**Author notes:** **Corresponding author; Present address:** Department of Microbiology and Parasitology, Faculty of Pharmacy, University of Medicine and Pharmacy at Ho Chi Minh City, Ho Chi Minh 700000, Vietnam. **Tu-Anh Nguyen E-mail address:**.

## Abstract

*Acinetobacter baumannii* is the leading cause of hospital-acquired infection in Vietnam. Of note, antibiotic resistance genes are significantly popular in clinical isolates of *A. baumannii*. Therefore, rapid identification of *A. baumannii* and determination of antibiotic resistance genes will help to make effective clinical decisions related to antibiotic use. This paper proposes a multiplex PCR to identify *Acinetobacter baumannii* and detect their ß-lactam antibiotic resistance genes in clinical isolates. Multiplex PCR was applied to amplified *rec*A gene and region ITS 16S - 23S rDNA for Rapid detection of *A. baumannii.* The two antibiotic resistance genes - *bla*_OXA-51-like_, *amp*C gene - were detected by multiplex PCR and three genes coding Extended-spectrum beta-lactamases - *bla*_*CTX-M*_, *bla*_*TEM*_, *bla*_*SHV*_ genes - were subjected to PCR. 49 bacteria strains were subjected to colony PCR. The result showed that 46 strains were *A. baumannii* and 3 strains belonged to the genus *Acinetobacter.* The multiplex PCR showed that all of 46 *A. baumannii* containing the *bla*_OXA-51-like_ gene and the *Amp*C gene; 34 strains possess the gene *bla*_TEM_ and none of them has *bla*_CTX-M_ and *bla*_SHV_ genes. The results of the multiplex PCR are the same as those of the *in vitro* antibiotic sensitivity testing of *A. baumannii*. However, applying the multiplex PCR directly from the bacteria colony, we can proceed simultaneously with the bacterial identification and the antibiotic resistance gene detection.

**Highlights:**

- 100% of isolates of *A. baumannii* contains the *bla*_OXA-51-like_ gene and the *Amp*C gene.
- 34/46 isolates possess the gene *bla*_TEM_, however, do not contain *bla*_CTX-M_ and *bla*_SHV_ genes.
- Combined disc test with cefotaxime/clavulanic acid/boronic acid is an excellent method to analyse ESBL phenotype.

## 1. Introduction

Currently, in the world as well as in Vietnam, *Acinetobacter baumannii* is one of the major causes of hospital – acquired infections [1]. Besides the increase in infection rate, there are *A. baumannii* strains possessing antibiotic resistance genes which code the enzyme hydrolyzing new generation antibiotics such as carbapenem. *A. baumannii* is also capable of producing the extended-spectrum beta-lactamase (ESBL) enzyme and thus can destroy most of the penicillin and broad-spectrum cephalosporin antibiotics [2]. According to a report by the Centers for Disease Control and Prevention (CDC) in 2013, there were about 63% of multidrug resistance *A. baumannii* infections [3]. Based on a review by the Global Antimicrobial Resistance Coordinator (2009), more than 60% of isolated *A. baumannii* were collected in some hospitals such as Bach Mai Hospital, Cho Ray Hospital and Central Hospital for Tropical Diseases (in Viet Nam) is multidrug-resistant [4].

For infectious diseases, rapid and accurate identification of pathogenic bacteria plays an important role in diagnosis and treatment. The traditional microbiological methods such as biochemical tests, antimicrobial susceptibility techniques, spend a lot of time. Meanwhile, applying the multiplex PCR, we can proceed simultaneously with the bacterial identification and antibiotic resistance gene detection. Therefore, research on the specific genes and antibiotic resistance genes in *A. baumannii* isolated from clinical samples by multiplex PCR was conducted. In this study, rapid detection of *A. baumannii* by multiplex PCR technique was carried out for determining the conservation gene *recA* that specified for the genus Acinetobacter and the region-specific ITS 16S - 23S rDNA of *A. baumannii* [5,6]. The antibiotic resistance genes including the *bla*_OXA-51-like_ gene - carbapenem resistance, *Amp*C gene - cephalosporin resistance [7,8] and three genes coding Extended-spectrum beta-lactamases (ESBL) - *bla*_*CTX-M*_, *bla*_*TEM*_, *bla*_*SHV*_ genes were selected [9–11]. There are four groups of enzymes in the OXA-type carbapenemase family that are commonly present in *A. baumannii*, OXA-23, OXA-24, OXA-58, OXA-51-like. Recently, scientists have discovered the *bla*_*Oxa*-51-like_ gene coding for the OXA-51-like enzymes on the chromosome and the natural antibiotic resistance gene of *A. baumannii*. Particularly, group A β-lactamase is the ESBL, it is able to destroy the third-generation cephalosporins such as ceftazidime, cefepime, cefotaxime and ceftriaxone, but not carbapenem. ESBLs mainly belong to the TEM and SHV β-lactamase groups, caused by mutations on one nucleotide of the *bla*_TEM_ and *bla*_SHV_ genes. In addition, ESBL types CTX-M, VEB-1, PER-1 are also common, especially ESBL CTX-M encoded by the *bla*_CTX-M_ gene.

## 2. Material and Methods

### 2.1. Bacteria strains

49 isolates of *Acinetobacter* spp., mostly from bronchial-fluid samples, were provided by the General Hospital in Binh Duong province, Viet Nam. These bacteria were identified by biochemical tests in the former experiments in this hospital. These isolates were cultured in Tryptic Soy Broth (TSB - Merck) before being streaked on MacConkey agar (MCA - Merck) to harvest colonies that were subjected to colony PCR.

### 2.2. Rapid detection of A. baumannii by multiplex PCR technique (RD-PCR)

The multiplex PCR technique (RD-PCR) was applied to determine the conservation gene *rec*A (425 bp) that specifies for the genus *Acinetobacter* with the primers P-rA1 - P-rA2 and the region-specific ITS 16S - 23S rDNA (208 bp) of *A. baumannii* with the primers P-Ab-ITSF - P-Ab-ITSB (**Table 1**) [5,6]. The RD-PCR reaction with a total reaction volume of a 25 μl consists of 2.5 μl PCR buffer (10X), 1.0 μl each of dNTPs (10 mM), 4.0 μl MgSO_4_ (25 mM), 1.0 μl each of primers (10 μM), 0.1 μl *Taq* DNA polymerase (5.0 UI), 1.0 μl bacterial suspension (one colony was dispersed in 20 μl TSB medium), and distilled water. PCR amplification process was carried out with an Labnet MultiGene™ OptiMax thermal cycler under the following conditions: initial denaturation (94 °C, 3 min); 30 cycles of denaturation (94 °C, 30 sec), annealing (51 °C, 30 sec), extension (72 °C, 45 sec); final extension (72°C, 5□min). PCR products were analysed by electrophoresis on 1% agarose gel (Sigma-Aldrich), containing SafeView (ABM), and UV visualization were performed on Gel Documentation System WGD-30 (DAIHAN Scientific Korean). *Bacillus subtilis* PY79 was used as a negative control.

**Table 1.**
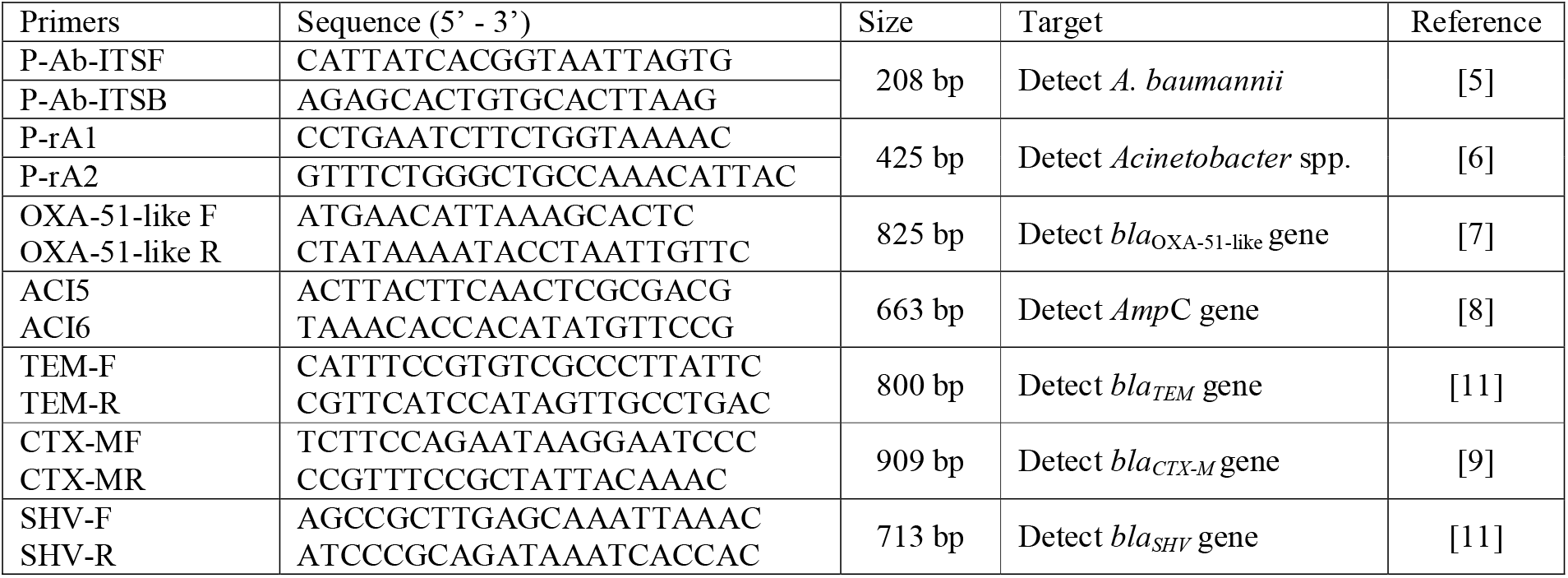
Primer sequences

### 2.3. Detection of the ß-lactamase genes of A. baumannii by multiplex PCR technique (B-PCR)

The B-PCR with a total reaction volume of a 25 μl includes 2.5 μl PCR buffer (10X), 1.0 μl each of dNTPs (10 mM), 4.0 μl MgSO_4_ (25 mM), 1.0 μl each of *bla*_OXA-51-like_ primers and ACI5 - ACI6 primers (Table 1) (10 μM), 0.5 μl *Taq* DNA polymerase (5.0 UI), 1.0 μl bacteria, and distilled water. The multiplex PCR program consists of initial denaturation (94 °C, 3 min); 30 cycles of denaturation (94 °C/ 1 min), annealing (47 °C /1 min), extension (72 °C /1 min); final extension (72 °C, 5□min). PCR products were detected by electrophoresis on 1% agarose gel [7,8]. Double distilled water was used as a negative control.

### 2.4. Detection of the ESBL genes of A. baumannii by multiplex PCR technique (E-PCR)

The E-PCR includes 2.5 μl PCR buffer (10X), 1.0 μl each of dNTPs (10 mM), 1.0 μl each of TEM, SHV, CTX-M primers (Table 1) and 0.2 μl *Taq* DNA polymerase (5.0 UI), 3.0 μl bacteria, and distilled water to adjust to a total volume of a 25 μl. The multiplex PCR program consists of initial denaturation (94 °C, 3 min); 30 cycles of denaturation (94 °C, 30 sec), annealing (55 °C, 1 min), extension (72 °C, 1 min); final extension (72 °C, 5□min). PCR products were detected by electrophoresis on 1% agarose gel [9–11]. Double distilled water was used as a negative control.

### 2.5. In-vitro antibiotic susceptibility testing

*A. baumannii* colonies were dispersed in 0.85% NaCl ~ 0.5 McFarland (1-3 ×10^8^ CFU/ml). Bacterial suspension of 10^6^ CFU/ml would be obtained by a 100 times dilution before being spreaded on Mueller Hinton Agar (MHA - Merck) [12]. The antibiotic discs, including imipenem (10 μg), cefepime (30 μg), cefotaxime (30 μg), amikacin (30 μg), colistin (10 μg), levofloxacin (5 μg), and doxycycline (30 μg) were used in disk diffusion method. These plates were incubated at 37°C for 24 hours in ambient air. Inhibitory zone diameters were measured by the electronic vernier caliper (Insize 1112–200). Paper discs were used as a negative control. The inhibitory zone was compared to CLSI M100-S28 [13].

### 2.6. Phenotypic detection of ESBL production by combined disc test

Plates were spreaded bacteria on the surface with the process being similar to section 2.5. For detecting ESBL producing isolates of *A. baumannii*, combination and alone antibiotic discs (Nam Khoa Biotek, Vietnam) were used in the disk diffusion test (Table 2) [14].

**Table 2.** Detection of ESBL production by combined disc test

| Method | Antibiotic | Testing medium | Interpretation of results | Ref. |
| --- | --- | --- | --- | --- |
| CEFOCLA | Using 2 antibiotic discs: cefotaxime (30 µg); cefotaxime/clavulanic acid (30/10 µg). | Muller Hinton Agar (MHA). | A ≥ 5 mm increase in inhibitory zone diameter (IZD) for a combination disc and a single antibiotic disc is regarded as ESBL positive (ESBL(+)), otherwise ESBL negative (ESBL(-)). | [15] |
| CEFOSUL | Using 2 antibiotic discs: cefotaxime (30 µg); cefotaxime/sulbactam (30/10 µg). | Muller Hinton Agar (MHA). | See above. | [15,16] |
| CLO-CEFOCLA | Using 2 antibiotic discs: cefotaxime (30 µg); cefotaxime/ clavulanic acid (30/10 µg). | MHA supplementary cloxacillin 200 µg/ml. | See above. | [17] |
| CEFO-CLA-BO | Using 4 antibiotic discs: cefotaxime (30 µg); cefotaxime/clavulanic acid (30/10 µg); cefotaxime/boronic acid (30/400 µg); cefotaxime/clavulanic acid/boronic acid (30/10/400 µg). | Muller Hinton Agar (MHA). | A ≥ 5 mm increase in IZD for a cefotaxime/clavulanic acid/boronic acid disc versus a cefotaxime disc or a ≥ 3 mm increase in IZD for a cefotaxime/clavulanic acid/boronic acid disc versus a cefotaxime/acid boronic disc indicates ESBL(+), otherwise ESBL(-). | [18,19] |

## 3. Results

### 3.1. Rapid detection of A. baumannii by multiplex PCR technique

There were 49 isolates of *Acinetobacter* spp. from the clinical samples that were used for multiplex PCR, and *Bacillus subtilis* PY79 was used as a negative control. The consequence is 46/49 bacterial isolates from the clinical samples were detected containing the two PCR products ~ 425 bp (*recA* gene) and ~ 208 bp (ITS 16S - 23S rDNA fragment), which are the specific nucleotide sequences of *A. baumannii* species. The only *recA* gene (425 bp) exists in the three other samples (5333, 4246, 5378) (**Figure 1**). This result indicated that 46 isolates are *A. baumannii* and 3 isolates belonged to the genus *Acinetobacter*. This identification result is the same as that of the biochemical method with commercial kit IVD NK-IDS 14 GNR (NamKhoa Biotek). Moreover, this experiment used the colony PCR technique that helps to shorten the time needed for DNA extraction, therefore it benefits from saving diagnostic costs.

**Figure 1.**
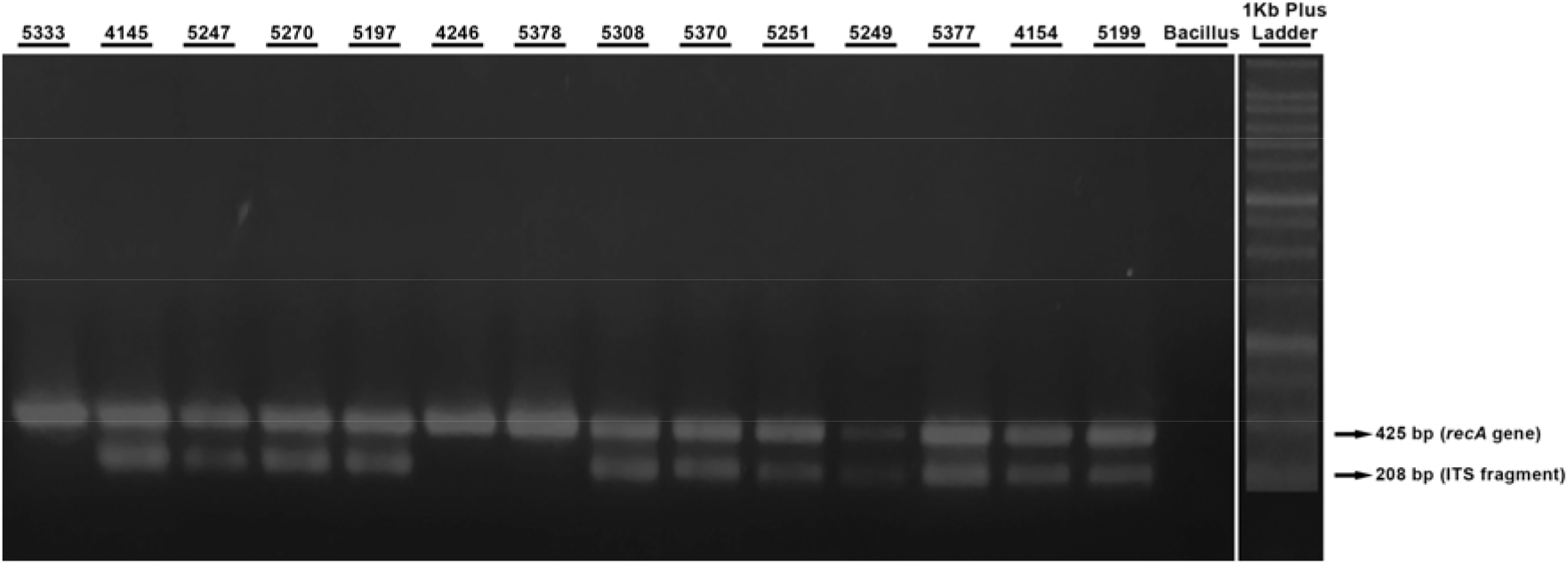
Multiplex PCR products for detection *A. baumannii;* B: *Bacillus subtilis* strain

**Figure 2.**
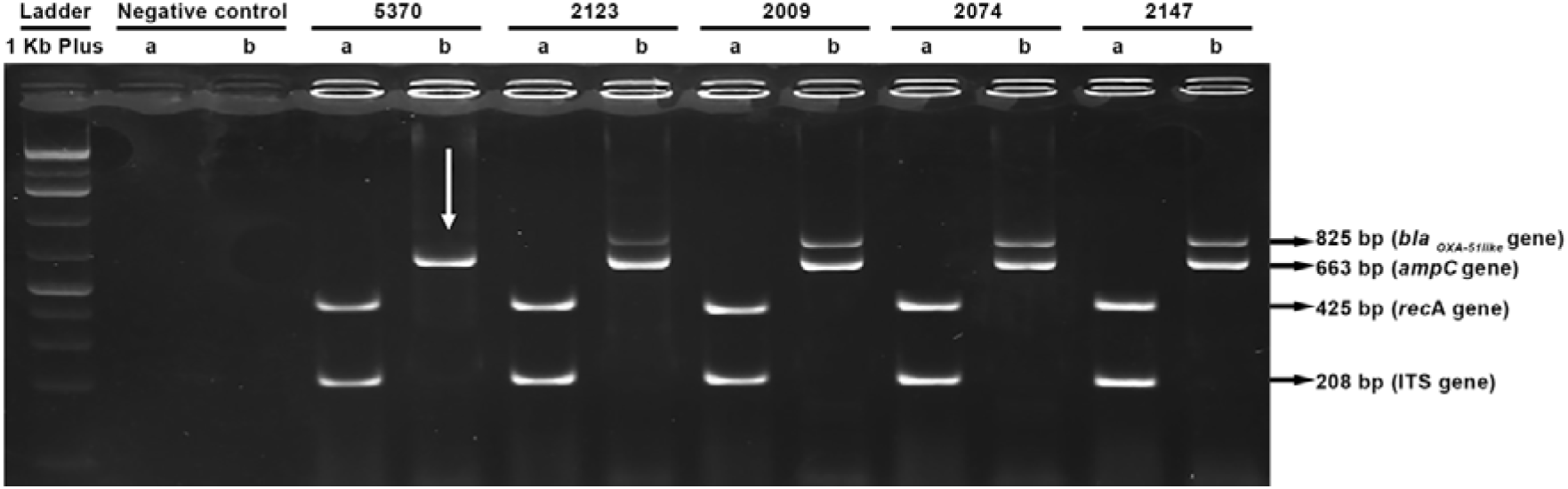
The specific PCR products of some *A. baumannii* isolates and their *ß-lactamase* genes (a) *recA* gene and ITS fragment; *(b) blaOXA-51* gene *and AmpC* gene. Sample 5370 contains only the *Amp*C gene.

**Figure 3.**
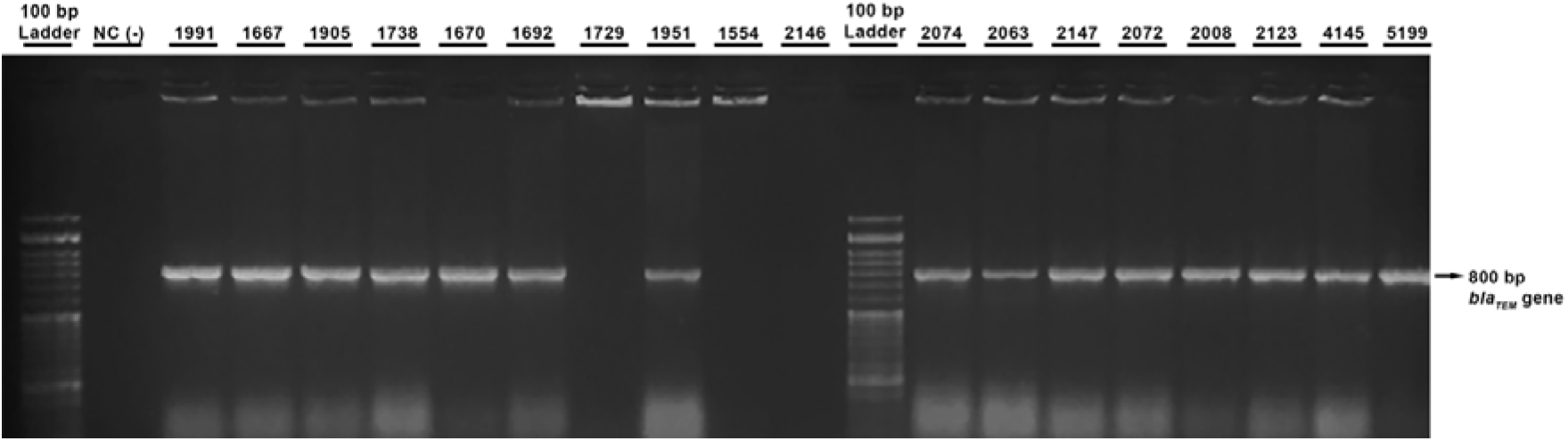
The specific PCR products of the *bla*_TEM_ gene. NC (−): negative control.

**Figure 4.**
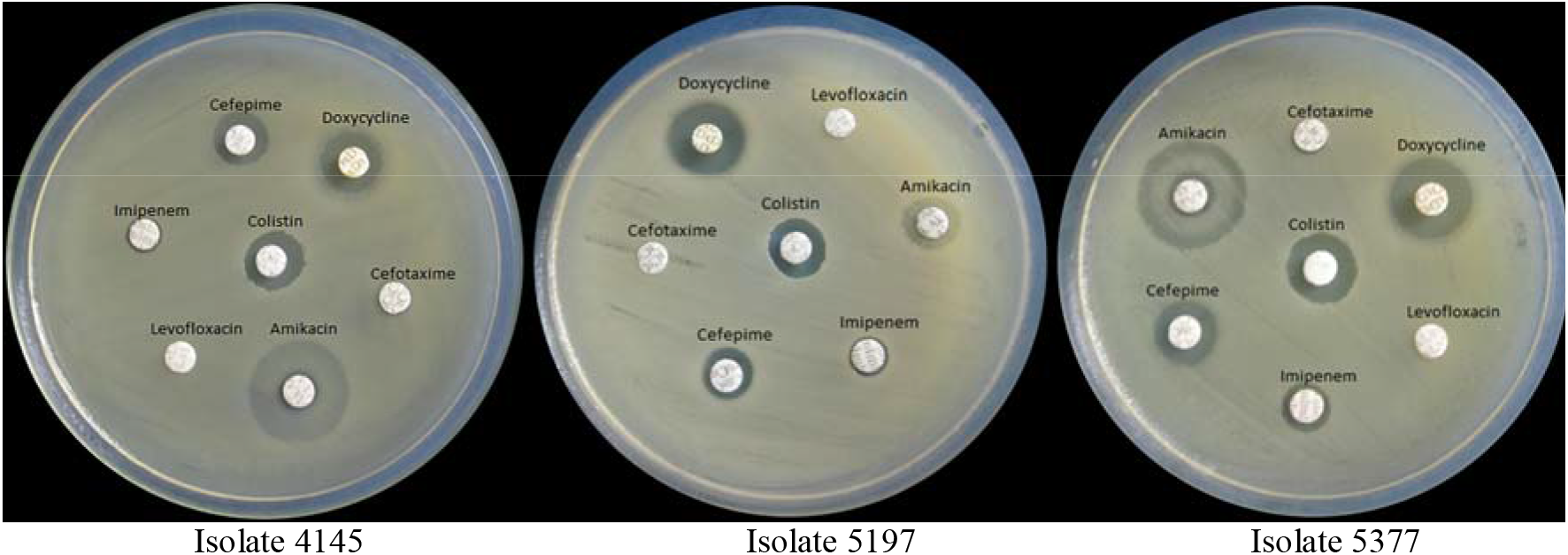
Antibiotic susceptibility testing of the three isolates 4145, 5197 and 5377

### 3.2. Detection of the ß-lactamase genes of A. baumannii

There are 45/46 *A. baumannii* isolates possessing two antibiotic resistance genes: *bla*_OXA-51-like_ ~ 825 bp (carbapenem resistance gene), *Amp*C ~ 663 bp (cephalosporin resistance gene), while the isolate 5370 expressed merely the *AmpC* gene. Therefore, the *Amp*C gene exists in all 46 isolates of *A. baumannii*; in other words, these isolates would be able to resist the antibiotic cephalosporin group. *A. baumannii* as well as some other Gram-negative bacteria are capable of producing the *AmpC* type β-lactamases by expression of *Amp*C gene on chromosome or plasmid. *AmpC* type β-lactamases can deplete penicillin, narrow and broad-spectrum cephalosporin, except carbapenem and fourth-generation cephalosporins such as cefepime, cefpirome [20]. 45 isolates displayed *bla*_OXA-51-like_ genes that encode for OXA-type carbapenemase [21]. This enzyme is responsible for hydrolysis of β-lactam antibiotics, especially carbapenem which is one of major antibiotics for treatment of severe infections caused by *A. baumannii*.

### 3.3. Detection of the ESBL genes of A. baumannii

34/46 isolates of *A. baumannii* contain *bla*_TEM_ gene (73.9%), however, none of the *bla*_CTX-M_ gene or *bla*_SHV_ gene was detected in all 46 isolates. This result seems to be extraordinary compared to other cases when *bla*_*CTX-M*_ gene is the most common among ESBL genes [10].

### 3.4. In vitro antibiotic susceptibility testing

Along with the detection of carbapenem, cephalosporin resistance genes by multiplex PCR, *in vitro* antibiotic susceptibility testing of 46 isolates of *A. baumannii* by antimicrobial susceptibility testing was also performed. Six employed antibiotic disc containing antibiotics those are usually used for the treatment of *A. baumannii* are imipenem, cefepime, cefotaxime, amikacin, colistin, levofloxacin, doxycycline. The inhibition zone of the 10 randomized representative samples (4145, 5199, 5308, 5249, 5370, 5247, 4154, 5197 and 5377) are shown in **Table 3** and **Figure 5** (all data shown in Supplementary 1).

**Figure 5.**
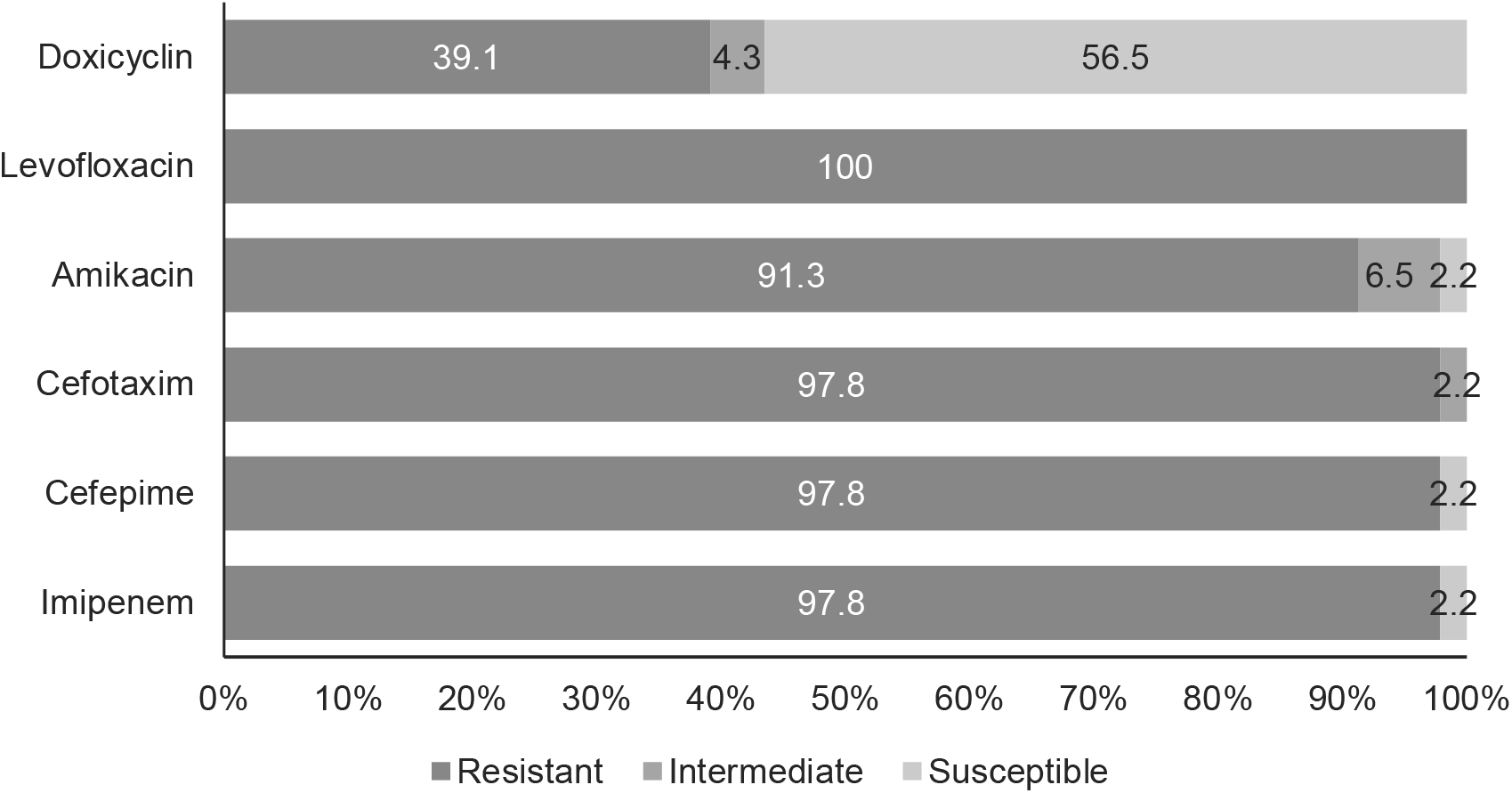
The rate of antibiotic resistance of 46 *Acinetobacter baumannii* isolates

**Table 3.** The inhibition zone of antibiotic susceptibility testing

| Isolates | Antibiotic |  |  |  |  |  |  |
| --- | --- | --- | --- | --- | --- | --- | --- |
|  | Im (mm)<br>(IZD/REF) | Cm (mm)<br>(IZD/REF) | Ct (mm)<br>(IZD/REF) | Ak (mm)<br>(IZD/REF) | Co (mm)*<br>(IZD/REF) | Lv (mm)<br>(IZD/REF) | Dx (mm)<br>(IZD/REF) |
| 4145 | R (9/≤18) | R (12/≤14) | R (7/≤14) | R (7/≤14) | S (12/≥11) | R (7/≤13) | S (15/≥13) |
| 5199 | R (8/≤18) | R (8/≤14) | R (7/≤14) | R (7/≤14) | S (11/≥11) | R (7/≤13) | S (15/≥13) |
| 5308 | R (8/≤18) | R (8/≤14) | R (7/≤14) | R (7/≤14) | S (11/≥11) | R (7/≤13) | S (14/≥13) |
| 5249 | R (8/≤18) | R (8/≤14) | R (7/≤14) | R (14/≤14) | S (11/≥11) | R (7/≤13) | S (16/≥13) |
| 5370 | S (28/≥22) | S (22/≥18) | I (18/15-22) | S (23/≥17) | S (11/≥11) | R (12/≤13) | S (13/≥13) |
| 5247 | R (8/≤18) | R (12/≤14) | R (7/≤14) | R (7/≤14) | S (12/≥11) | R (7/≤13) | S (16/≥13) |
| 5270 | R (10/≤18) | R (10/≤14) | R (7/≤14) | I (15/15-16) | S (14/≥11) | R (7/≤13) | S (19/≥13) |
| 4154 | R (8/≤18) | R (11/≤14) | R (7/≤14) | R (7/≤14) | S (11/≥11) | R (7/≤13) | S (14/≥13) |
| 5197 | R (9/≤18) | R (10/≤14) | R (7/≤14) | R (7/≤14) | S (11/≥11) | R (7/≤13) | S (15/≥13) |
| 5377 | R (7/≤18) | R (9/≤14) | R (7/≤14) | R (7/≤14) | S (12/≥11) | R (7/≤13) | S (15/≥13) |

IZD: inhibitory zone diameter of antibiotic resistance testing, REF: inhibitory zone diameter of antibiotic sensitivity to *A. baumannii* according to CLSI. Im: Imipenem, Cm: Cefepime, Ct: Cefotaxime, Ak: Amikacin, Co: Colistin, Lv: Levofloxacin, Do: Doxycycline. R: Resistance; S: Sensitivity; I: Intermediate resistance. \* In this research, we used colistin discs as a preliminary test, however, updated CLSI's guideline [13] recommends using MIC value for colistin because of its' poor diffusion. Therefore, we did not discuss this based on the IZD of colistin.

### 3.5. Phenotypic detection of ESBL production by combined disc test

The method CEFOCLA used cefotaxime alone and combined cefotaxime/clavulanic acid, this test displayed ESBL(+) on two isolates, 2146 and 1672. Meanwhile, almost all isolates show no inhibitory zone with combined discs containing cefotaxime/sulbactam, hence, CEFOSUL is not suitable for identification of ESBL phenotype. We recorded 7 isolates ESBL(+) including 5377, 5197, 5249, 5199, 2146, 1692, 1672 when using MHA containing cloxacillin 200 μg/ml for combined disc test with cefotaxime and cefotaxime/clavulanic acid. Cloxacillin is known as an AmpC β-lactamase inhibitor [17].

Based on the CEFO-CLA-BO method, we identified 40 out of 46 isolates (87%) expressing ESBL(+) in which there are 34 isolates possessing *bla*_*TEM*_ and there is not isolate displaying *bla*_*SHV*_ and *bla*_*CTX-M*_. The AmpC gene presents in all isolates of this research. This gene encodes for AmpC β-lactamase which is not inhibited by clavulanic acid or sulbactam. Therefore, the CEFO-CLA-BO method is effective in applying boronic acid to inactivate the AmpC β-lactamase enzyme [18].

## 4. Discussion

Only isolate 5370 sensitive to the carbapenem group (imipenem), the remaining 45 out of 46 isolates were resistant to this antibiotic (97.8%). This result is reasonable because no detection of the *bla*_OXA-like-51_ gene in isolate 5370 as shown above. The same as the case of the fourth-generation cephalosporin (cefepime), only the isolate 5370 is sensitive and the other 45 isolates are resistant (97.8%), although all 46 isolates contain *Amp*C genes. To be able to discuss this phenomenon, the *Amp*C gene of the 5370 should be sequenced for comparing with other *Amp*C gene sequences. Perez F. et al. (2007) showed that although the bacteria have the *Amp*C gene, they resist to cephalosporins except for cefepime [20]. With the third-generation cephalosporin (cefotaxime) the resistance and intermediate resistance rate is up to 100%. This is similar to the multiplex PCR results when all of the tested *A. baumannii* have the cephalosporin resistance *Amp*C gene. About the other antibiotic susceptibility, the aminoglycoside group (amikacin) has a high rate of resistance and intermediate resistance (97.8%). For the fluoroquinolone group, *A. baumannii* is 100% resistance to levofloxacin. The cycline group (doxycycline), bacterial sensitivity is about 56.5%, the rate of resistance and intermediate resistance about 43.5%.

This study shows that *A. baumannii* isolated in Binh Duong hospital have been resistant to many groups of antibiotics with high rates. This result is similar to the reported article which mentioned antibiotic resistance of *A. baumannii* by the Pasteur Institute in Ho Chi Minh City, Viet Nam [22]. As reported by the Pasteur Institute [22], the percentage of cephalosporin and amikacin resistance was 96.7%, while in this study, the rate of resistance cephalosporin was 100%, of amikacin was 97.8% (based on antibiotic susceptibility testing). Noticeably, the carbapenem group was used to be the priority choice antibiotics for infection caused by multidrug resistance *A. baumannii*, but the high rate of resistance was reported 96.7% and 97.8%, respectively according to the Pasteur Institute [22] and this study.

By multiplex PCR, we identified 46 *A. baumannii* isolated from different clinical departments, which were resistant to cephalosporins. 45 of 46 samples carry the *bla*_OXA-51-like_ gene which is a specific gene of *A. baumannii*. Among the 46 strains of *A. baumannii* which resist third-generation cephalosporins, we detected the *Amp*C gene. The *Amp*C gene located on the chromosome or plasmid and encodes the class C β-lactamase - AmpC cephalosporinase enzyme [23].

Due to the existence of the *bla*_TEM_ gene in the 34 strains of *A. baumannii*, this is evidence of their ability to produce ESBL. Most data of the ESBL molecular method align with the phenotypic test, however, 6 isolates (1571, 1672, 1793, 1958, 2146, 5251) expressed ESBL(+) in disc test only. ESBL(+) associates with different mechanisms, in this research we targeted 3 popular genes encoded for ESBL including *bla*_SHV_, *bla*_CTX-M_, *bla*_TEM_. This phenomenon could be due to *bla*_*SHV*_ and *bla*_*CTX-M*_ being not popular in Binh Duong area, while *bla*_TEM_ displays in 73.9% isolates of *A. baumannii* in this study.

Based on the previous researches, the type of genes responsible for extended-spectrum β-lactamases is different in local areas. According to a report in Saudi Arabia in 2015 [10], there were 71% *bla*_TEM_ gene, none of *bla*_SHV_ gene and 81% *bla*_CTX-M_ gene. While research of Safari M et al. (2015) in Hamadan City in Iran [9] showed that there were *bla*_CTX-M_ 20%, *bla*_SHV_ 58% and *bla*_TEM_ 0% from the isolated *A. baumannii*. However, following the research on *A. baumannii* from Tehran hospitals of Melika Sharif *et al*, these rates were *bla*_SHV_ 63% and *bla*_TEM_ 56% [11].

## 5. Conclusions

Acquired hospital infection caused by the Gram (−) bacteria of the genus *Acinetobacter* is a serious problem due to increasing the severity of the disease, lengthening treatment duration, increasing treatment costs and increasing mortality. Thus, the rapid and accurate identification of *A. baumannii* as well as their resistance antibiotic genes plays an important role. Applying the molecular method can overcome these above difficulties. Particularly, the colony PCR technique helps not only to shorten the consumption of time needed for bacterial culture and the extraction of chromosomal DNA but also to save on diagnostic costs. Furthermore, based on the rapid result of multiplex PCR done directly from colony, clinicians can prescribe effective antibiotics, resulting in reduction of the incidence of this disease as well as patients will also progress better. Meanwhile, identification of bacteria or of their antibiotic susceptibility by traditional microbiology methods such as biochemical tests and antimicrobial susceptibility tests take a lot of time, effort and in some cases it is difficult to explain unclear identification results.

Two multiplex PCR detection procedures were set up including detection of the specific genes of the strain *A. baumannii* based on the *rec*A gene and the ITS 16S - 23S rDNA fragment, ß-lactamase producers based on the two genes *bla*_OXA-51_ gene coding carbapenemase and the *Amp*C gene coding cephalosporinase. In addition, one simple PCR test were designed for detecting *bla*_TEM_ gene. The results of this tests are the same as those of the *in vitro* antibiotic sensitivity testing of *A. baumannii* using the antimicrobial susceptibility technique except for the case of screening ESBL using the combination disc method.

## 6. Acknowledgements

We would like to thank General Hospital Binh Duong for providing the *A. baumannii* from the clinical isolates.

Ethical Approval

